# A multilevel hierarchical framework for quantification of experimental heterogeneity

**DOI:** 10.64898/2025.12.21.695338

**Authors:** David J. Warne, Xiangrun Zhu, Thomas P. Steele, Stuart T. Johnston, Scott A. Sisson, Matthew Faria, Ryan J. Murphy, Alexander P. Browning

## Abstract

Biological systems exhibit substantial heterogeneity: that is, variation in specific characteristics of individuals within a population. As a result, it is of critical importance to appropriately account for biological heterogeneity when calibrating mathematical models to infer cellular processes and predict behaviour. Recent approaches consider ordinary differential equations with random parameters to quantify heterogeneity in dynamical processes of cells. In this setting, statistical inference is performed to characterise the distribution of these random parameters within a cell population. One significant limitation of this approach is the tacit assumption that there are no substantial deviations in these distributions across experimental replicates. In this work, we propose a flexible Bayesian hierarchical differential equation modelling framework that quantifies and distinguishes both inter-experimental heterogeneity (heterogeneity between experimental replicates) and intra-experimental heterogeneity (biological heterogeneity within replicate populations). We consider two recent studies that employ mathematical models to interpret flow cytometry snap-shot data and quantify heterogeneity in nano-particle cell interactions and cell internalisation processes. Using simulation data, we demonstrate that substantial inaccuracy in the inferred dynamics can arise when experimental heterogeneity is not accounted for. By contrast, our hierarchical approach is robust to variability in inter-experimental and intra-experimental heterogeneity and our method simplifies to previous methods when inter-experimental heterogeneity is negligible. Our approach is flexible and widely applicable to applications involving replicate populations and snapshot data.

## 1 Introduction

Accounting for biological heterogeneity is crucial in the biomedical sciences [1]. Heterogeneity plays a key role in the development and maintenance of healthy living organisms [2, 3] through complex interactions between various types of cells driven by intracellular biochemical processes [4]. In pathological situations, such as cancer or viral infections [5–8], heterogeneity can seriously impact the efficacy of pharmaceutical interventions and eventual patient outcomes [9–11]. As a result, appropriately quantifying biological heterogeneity within a population of cells is of tremendous importance for understanding natural processes and for the design of effective drugs, especially in the areas of precision medicine such as targeted therapeutics [12–15].

A standard experimental approach to quantify properties of cell populations is flow cytometry [16–18]. In such an experiment, populations of cells (Figure 1(a)), are tagged or stained with fluorescent markers. The cells then individually flow through the cytometer nozzle and pass through a laser from which scattered light is amplified and converted to a digital signal representing a measurement of fluorescent intensity related to each cell in the population (Figure 1(b)). This results in a snapshot in time of the fluorescent intensity distribution across multiple colour channels, providing a rich source of information about potential variability in the cell populations at a single point in time. (Figure 1(c)). Typically, thousands to tens of thousands of cells are analysed per time point.

**Figure 1:**
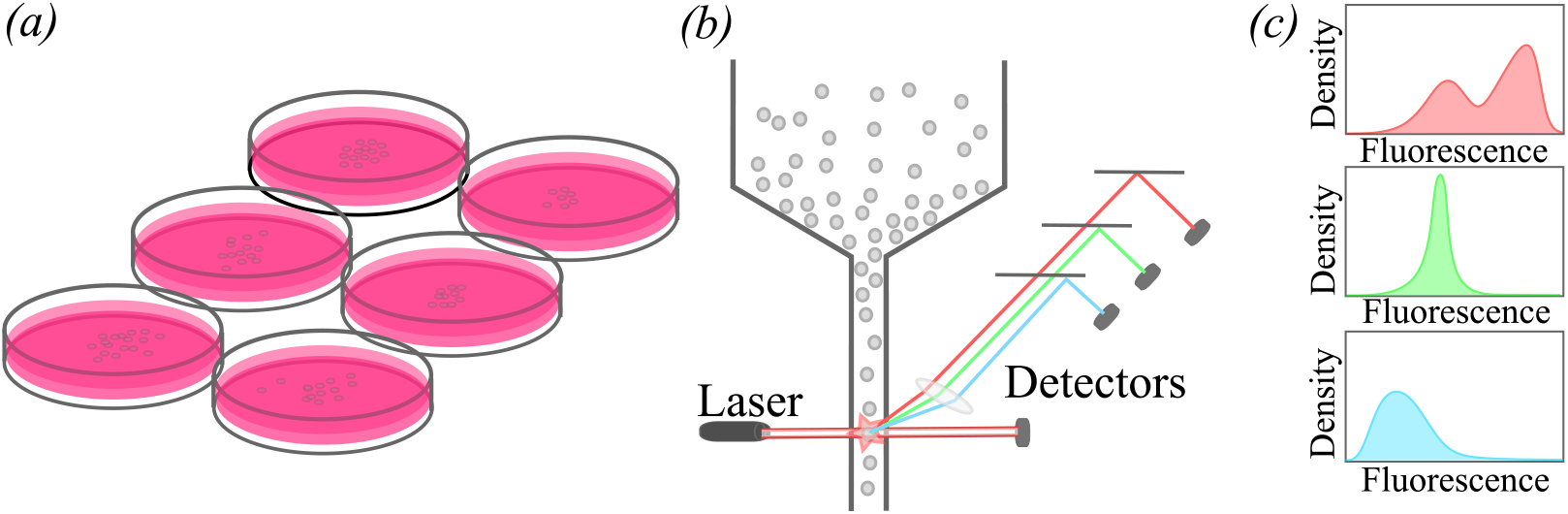
Schematic of flow cytometry data acquisition. (a) Cells are seeded into cell culture plates e.g., 12-well plates, in a growth media (e.g., 1 mL) and incubated with various treatments for some time period (typically between 1 to 48 hours). (b) Cells are then removed from plates and placed in suspension, and passed through the flow cytometer nozzle to a laser beam where light scattering from the cell is measured at detectors. (c) For each fluorescent channel, the intensity distribution is obtained, leading to distributions showing cellular variability.

Data obtained from flow cytometry experiments exhibit substantial levels of heterogeneity. While some of this variation can be attributed to measurement noise, recent studies have demonstrated that some of this variation arises due to intrinsic biological variation between cells despite them being genetically identical [19, 20]. However, there are also extrinsic factors that drive heterogeneity in flow cytometry experiments that can negatively affect the statistical analysis of the biological heterogeneity of interest if not properly accounted for [21, 22]. For example, subtle variation in sample preparation, instrument configuration for data acquisition, and manual user operation can have a substantial effect on the signal-to-noise ratio and statistical bias in the data, potentially impacting the validity of conclusions drawn from the analysis [21–24].

In many studies, it is typically assumed that experimental replicates of identically prepared cell populations represent independent and identically distributed replicates. For example, in the work of Faria et al. [25], human leukaemia cells (THP-1 cell line) are seeded into a 12-well plate and nanoparticle-cell interactions are observed via flow cytometry at six observation times (between 1 hr and 24 hrs), resulting in two replicates per observation time. In practice, statistical analysis is performed on the pooled sample of replicates [26, 27], or each replicate is individually analysed [28]. Neither approach is ideal. Pooling ignores heterogeneity between experimental replicates and may lead to incorrect analysis.

Conversely, analysing replicates independently can lead to difficulties in assessing global trends [29]. A potential solution to analyse flow cytometry data is to use mathematical or statistical models that structure relationships between snapshots, and between time points data [29–31].

In this work, we introduce an approximate Bayesian multilevel hierarchical modelling framework for quantifying biological heterogeneity in flow cytometry data in situations where the parameters of interest are related to a mathematical model of cell dynamics. To demonstrate our approach, we consider simulation studies based on two recent studies that quantify biological heterogeneity in cell behaviour using pooled flow cytometry data [19, 32]. We highlight the dramatic impact that variation between replicates can have on pooled sample analysis results and show how a hierarchical approach can account for this effect. In addition, our approach reduces to the pooled case in situations where variation between replicates is minimal. Our approach is therefore generally applicable for applications in which controlling variation across flow cytometry replicates is infeasible.

## 2 Materials and Methods

We develop a general Bayesian hierarchical modelling framework for the analysis of heterogeneity in cell behaviour using flow cytometry data. This heterogeneity is variability that can be intra-experimental heterogeneity within a replicate, or inter-experimental heterogeneity between replicates. This inter-experimental heterogeneity can be variation between either technical replicates or experimental replicates. For clarity, we define a technical replicate to be an identically prepared replicate performed at the same time and the same person, and define an experimental replicate to be a replicate replicate performed on a different day, and/or by a different person (See Figure 2). For example, if an experiment was performed with a 12-well plate where two-wells were assigned to each of six time points then this would represent two technical replicates. However, if the entire 12-well plate experiment was performed again at a different lab or with a different instrument then this would be two experimental replicates, each with two technical replicates. For most experimental protocols, we expect technical replicates to be closely related. As a result, we focus our attention to experimental replicates in this manuscript. However, we show how this can be extended to account for both technical and experimental replicates in Appendix A.

**Figure 2:**
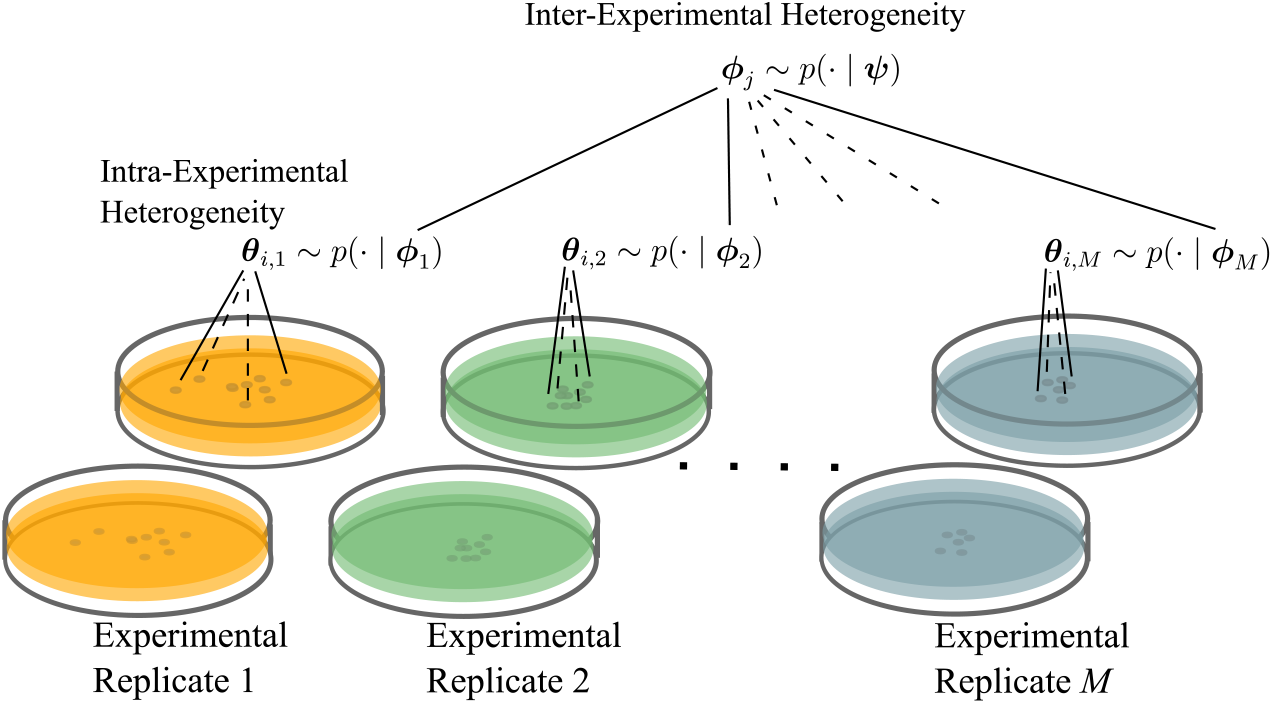
A schematic representation of the hierarchical statistical framework. Experimental replicates are indicated by different colours, with each experiment consisting of two identical technical replicates. Inter-experimental heterogeneity is captured through the hyper-parameters ***ψ*** that characterise the distribution of the *M* replicate specific hyper-parameters, ***ϕ***_1_, ***ϕ***_2_, …, ***ϕ***_*M*_. Here, ***ϕ***_*j*_ characterises the intra-experimental heterogeneity between *N* cells with ***θ***_*i,j*_ being the dynamic parameters for the *i*th cell within the *j*th replicate.

### 2.1 Flow cytometry data

We use simulated flow cytometry data that is a realistic characterisation of real flow cytometry data. Specifically, we build our data simulation process based on recent studies that investigate heterogeneity in interactions between cells and nano-particles [25, 26, 32] and cell internalisation processes [19, 33, 34].

#### 2.1.1 Nano-engineered particle-cell interaction data

The flow cytometry data published by Faria et al., [25] and subsequently analysed for heterogeneity in particle-cell interaction by Murphy et al., [32] are for a human leukaemia monocytic suspension cell line, THP-1. Here, THP-1 cells are seeded in a 12-well plate and incubated with nano-particles (specifically, we focus on the 150 nm core-shell data) for 1, 2, 4, 8, 16, and 24 [hr]. After incubation, nano-particles that are not associated with cells are removed via washing. The nano-particle fluorescence across the cell population from each well plate is collected via flow cytometry. This results in snapshot data of associated nano-particle fluorescence for *N* ≈ 20, 000 cells for each incubation time point.

#### 2.1.2 Dual-labelled probe internalisation data

Browning et al., [19] published flow cytometry data for a human B Cell lymphoblast cell line, C1R cells, and analysed the heterogeneity in endocytosis of anti-transferrin (anti-TFR) antibodies. The measurement technique is based upon programmable sensors using DNA quenching probes [35, 36]. Antibodies are dual-labelled with fluorescent dyes: FIP-Cy5 and BODIPY FL. Of these two probes, FIP-Cy5 is quenchable, that is, the fluorescence is effectively disabled on the cell surface using a quencher dye. Here, C1R cells are incubated with dual-labelled antibodies with incubation times of 5, 10, 20, 30, 60, 120, and 180 [min]. After incubation, cells are washed and resuspended both with and without the quencher [36]. The fluorescent signals for both FIP-Cy5 and and BODIPY FL are measured for the cell populations in both quenched and unquenched samples via flow cytometry. This leads to snapshots of the jointly distributed FIP-Cy5 and BODIPY FL fluorescence for *N* = 1, 000 cells over the incubation times and qenched/unquenched samples.

### 2.2 Mathematical modelling

We consider applications for which flow cytometry data is analysed using deterministic modelling based on ordinary differential equations (ODEs) as is commonplace in the mathematical and computational biology communities. The goal of such analysis is to quantify biological heterogeneity in parameters related to the dynamics of some cellular process. In this sense, each individual cell in the population is treated as having its own set of parameters and the goal is to find a suitable distribution for the parameters leading to cell dynamics matching the data distribution over time [19, 32, 37]. Here we describe the general ODE framework with random parameters used in the setting of pooled replicates, and then connect the framework to the particle-cell interaction model of Murphy et al., [32] and the cell internalisation model of Browning et al., [19]. These previous works focus exclusively on intra-experimental heterogeneity. In Section 2.3, we then show how to extend the analysis framework also account for inter-experimental heterogeneity between replicates and present a Bayesian multilevel hierarchical approach to quantify both intra- and inter-experimental heterogeneity together.

#### 2.2.1 Cell population models with random parameters

We consider a heterogeneous population of *N* cells. Let **X**_*i*_ be a vector of *d*_*X*_ *≥* 1 dynamic properties of interest related to the *i*th cell (e.g., the average number of nano-particles associated with cell, or concentrations of various biochemicals). This property vector evolves over time according to

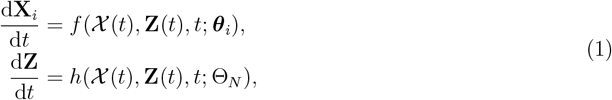

for *i* = 1, 2, …, *N*, where ***θ***_*i*_ is a vector of *d*_*θ*_ ≥ 1 parameters with values specific to the *i*th cell, Θ_*N*_ = [***θ***_1_, ***θ***_2_, …, ***θ***_*N*_] is a *d*_*θ*_*×N* matrix of the populations parameters, 𝒳(*t*) = [**X**_1_(*t*), **X**_2_(*t*), …, **X**_*N*_ (*t*)] is a *d*_*X*_ × *N* matrix representing the state of the cell population, and **Z**(*t*) is a vector of *d*_*Z*_ *≥* 1 shared environmental variables that interacts with cells in the population (e.g., a shared nutrient source). The functions *f* (*𝒳* (*t*), **Z**(*t*), *t*; ***θ***_*i*_) and *h*(*𝒳* (*t*), **Z**(*t*), *t*; Θ_*N*_) define the dynamics of cells and the environment, respectively. For *t*_0_ representing the start of the initial incubation time, we have initial conditions **X**_*i*_(*t*_0_) = **x**_*i*,0_ for *i* = 1, 2, …, *N*, and **Z**(*t*_0_) = **z**_0_. Importantly, we do not assume that these initial conditions are known quantities, since such assumptions are known to cause problems in study reproducibility [38]. Instead, initial conditions are included as model parameters that we will estimate.

The measured fluorescence of each cell is subject to external noise and thus treated as a random variable, that is,

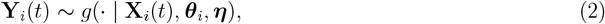

where **Y**_*i*_(*t*) are the recorded fluorescent intensities for the *i*th cell given the cell state **X**_*i*_(*t*), dynamics parameters ***θ***_*i*_, and observation process specific parameters, ***η***. The functional from of the observation process, *g*(· | **X**_*i*_(*t*), ***θ***_*i*_, ***η***), will typically be determined based on calibration data for flow cytometry. The dimension of **Y**_*i*_(*t*) will be the number of fluorescence channels and will generally not be the same as the dimension of **X**_*i*_(*t*). Thus a single cell population snapshot obtained at time *t > t*_0_ using flow cytometry is given by *𝒴*(*t*) = [**Y**_1_(*t*), **Y**_2_(*t*), …, **Y**_*N*_ (*t*)]. Typically, we do not observe the initial condition at *t* = *t*_0_, and therefor it is treated as a latent variable to infer with the model parameters. Finally, for *n* observation times, *t*_1_ *< t*_2_ *<* … *< t*_*n*_, we represent the entire flow cytometry dataset as 𝒟= [𝒴 (*t*_1_), *𝒴* (*t*_2_), …, *𝒴* (*t*_*n*_)].

Since snapshot data as obtained through flow cytometry does not track individual cells, using such data to perform parameter inference on ***θ***_1_, ***θ***_2_, …, ***θ***_*N*_ is not meaningful. Instead, one can consider ***θ***_1_, ***θ***_2_, …, ***θ***_*N*_ to be independent identically distributed (i.i.d.) samples from a parametric distribution with probability density, *p*(***θ*** | ***ϕ***), then perform parameter inference on the hyper-parameters ***ϕ*** ∈ **Φ**, where **Φ** is the hyper-parameter space. In a Bayesian setting, this leads to

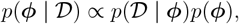

where

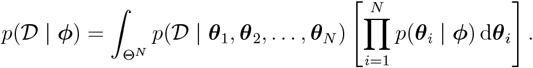

This is the essence of the approach taken by Browning et al., [19] and Murphy et al.,[32] and it relies on pooled snapshot data that requires the assumption of i.i.d. parameters across all cells and replicates. Our main contribution, described in Section 2.3, is an extension to this framework that only requires i.i.d. parameters within each experimental replicate.

#### 2.2.2 Example 1: Particle-cell interaction model

The particle-cell interaction model of Murphy et al., [32], describes the association of free nano-particles to a population of cells. Given *N* cells the number of particles associated with the *i*th cell at time *t > t*_0_, *P*_*i*_(*t*), is governed by the system,

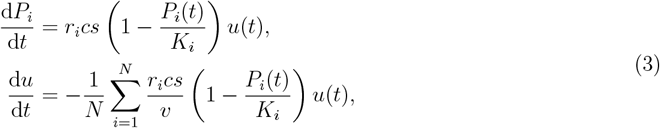

for *i* = 1, 2, …, *N*, where *r*_*i*_ and *K*_*i*_ are, respectively, the association rate and carrying capacity of particles for the *i*th cell, *c* is the fractional cell surface coverage, *s* is the surface area of the cell boundary, *v* is the volume of the well-mixed media, and *u*(*t*) is the total free particle density with initial condition *u*(*t*_0_) = *u*_0_. In the context of our framework given in Section 2.2.1, we have **X**_*i*_(*t*) = *P*_*i*_(*t*) and **Z**_*i*_(*t*) = *u*(*t*) (i.e., *d*_*X*_ = *d*_*Z*_ = 1), and ***θ***_*i*_ = [*r*_*i*_, *K*_*i*_]^⊤^ (i.e., *d*_*θ*_ = 2).

The measured fluorescence, *Y*_*i*_(*t*), for cell *i* is modelled by

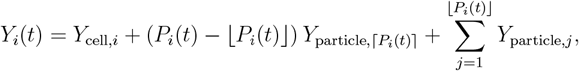

where *Y*_cell,*i*_ ∼ *p*(· | ***η***) is the autofluorescence of a cell, drawn at random from the empirical distribution cell-only calibration dataset, and 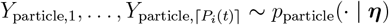 are individual particle fluorescences drawn at random from the empirical distribution particle-only calibration dataset. Here ***η*** are parameters associated with normalising voltages that are used to obtain flow cytometry measurements (See Murphy et al., [32] for details). Since there are *N* = 20,000 cells, this leads to a computationally challenging system of ODEs as they are fully coupled through the environment (Equation (3)). However, a tractable approximation with a semi-closed form solution can be obtained under the assumption that *u*(*t*) ≈ *u*_0_ throughout the simulation time (See Murphy et al., [32] for details). This assumption is quite common in the field and represents a high nano-particle dose such that the cells are not expected to meaningfully deplete it. Importantly, it does not lead to a completely decoupled system free from any environmental coupling and changes in *u*(*t*) are still captured.

#### 2.2.3 Example 2: Internalisation model

Browning et al., [19] consider a model describing the internalisation of transferrin anti-bodies that accounts for receptor recycling. Due to the experimental protocol (Section 2.1.2), the concentration of free transferrin antibodies can be considered sufficiently high that unbound surface receptors are assumed to bind instantaneously to a free antibody. Furthermore, the high free antibody concentration enables the internalisation dynamics of all *N* = 2,000 cells to be considered independent of each other since cells will never compete for free antibodies. For the *i*th cell at time *t > t*_0_, Browning et al., [19] model the dynamics of the density of unbound internal receptors, *T*_*i*_(*t*), bound surface receptors, *S*_*i*_(*t*), internalised bound receptors, *E*_*i*_(*t*), and interalised free antibodies, *F*_*i*_(*t*), according to,

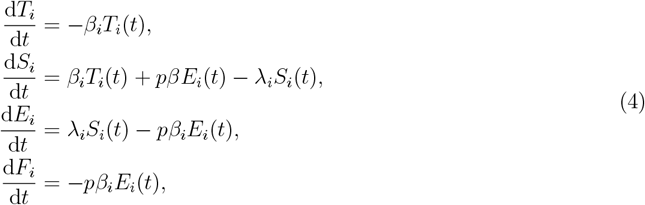

for *i* = 1, 2 …, *N*. Here, the initial total density of receptors for the *i*th cell is given by *T*_*i*_(*t*_0_) + *S*_*i*_(*t*_0_) = *R*_*i*_ and it is assumed no antibodies have initially been internalised with *E*_*i*_(*t*_0_) = *F*_*i*_(*t*_0_) = 0. Further, *λ*_*i*_ *>* 0 and *β*_*i*_ *>* 0 are, respectively, the internalisation rate and recycling rate for the the *i*th cell. Finally, *p* ∈ (0, 1] is the disassociation probability that captures the probability that an internalised antibody is absorbed into the cell versus being recycled back to the surface. Following Browning et al., [19], *p* is associated with a purely chemical process and considered constant across cells, whereas *λ*_*i*_, *η*_*i*_ and *R*_*i*_ are assumed to be heterogeneous across cells *i* = 1, 2, … *N*. Relating this to our framework (Section 2.2.1), we have **X**_*i*_(*t*) = [*T*_*i*_(*t*), *S*_*i*_(*t*), *E*_*i*_(*t*), *F*_*i*_(*t*)]^⊤^ (i.e., *d*_*X*_ = 4), and ***θ***_*i*_ = [*λ*_*i*_, *β*_*i*_, *R*_*i*_, *p*]^⊤^ (i.e., *d*_*θ*_ = 4). In this model we do not have a dependent shared environment so *d*_*Z*_ = 0. Due to the independence assumption, an analytic solution to Equation (4) can be obtained using the matrix exponential (See Browning et al., [19] for details).

The observation process is more complex in this model than the particle-cell interaction model (Section 2.2.2). Here, the experimental protocol involves dual-labelled antibodies with one of the labels (FIP-Cy5) being “quenchable”, leading to a loss in FIP-Cy5 fluorescence for free or surface bound antibodies, and the other (BODIPY FL) being “unquenchable” with fluorescence unaffected by the quenching process. The *N* = 2, 000 cells consist of two equal-sized technical replicates of size 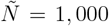 with one replicate undergoing the *quenching* process before being processed through the flow cytometer. This leads to four measured fluorescence signal values for the *i*th pair of cells,

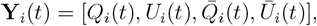

where *Q*_*i*_(*t*) (resp. *U*_*i*_(*t*)) are the quenchable FIP-Cy5 (resp. unquenchable BODIPY FL) fluorescence signals measured from the unquenched sample group, and 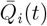 (resp. 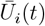) are the quenchable FIP-Cy5 (resp. unquenchable BODIPY FL) fluorescence signals measured from the quenched sample group. The resulting observation process is

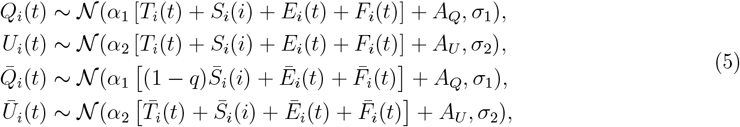

for 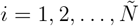, where *α*_1_ and *σ*_1_ (resp. *α*_2_ and *σ*_2_) are intensity and noise the fluorescence measurement for FIP-Cy5 (resp. BODIPY FL). Further, *A*_*Q*_ (resp. *A*_*U*_) corresponds to the average autofluorescence of FIP-Cy5 (resp. BODIPY FL), and *q* is the quenching efficiency. While *A*_*Q*_, *A*_*U*_ and *q* are pre-estimated from data directly, the parameters ***η*** = (*α*_1_, *α*_2_, *σ*_1_, *σ*_2_) must be inferred with the model parameters.

### 2.3 Statistical framework

In Section 2.2.1 we describe the typical setting in which biological heterogeneity has been treated in the literature. That is, replicates are pooled and treated as a single snapshot population [19, 25–27, 32]. In this work, we propose an extension in which each replicate is treated as its own population with variation within replicates (i.e., intra-experimental heterogeneity) being distinguished from variation between replicates (i.e., inter-experimental heterogeneity). To simplify the exposition of our approach we present a two-level Bayesian hierarchical model, however, the approach can be extended to more levels as shown in the Appendix A.

#### 2.3.1 Inter-experimental and intra-experimental heterogeneity

Within the random ODE framework given in Section 2.2.1, we consider *M* experimental replicates each containing *N* cells in total across identical technical replicates (Figure 2). The cell population dynamic model (Equation (1)) becomes,

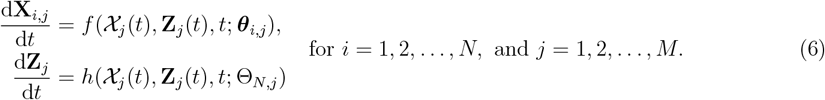

Here, **X**_*i,j*_(*t*) is the state of the *i*th cell in the *j*th replicate having parameters ***θ***_*i,j*_. For the *j*th replicate we have the subpopulation state, *𝒳*_*j*_(*t*) = [**X**_1,*j*_(*t*), **X**_2,*j*_(*t*), …, **X**_*N,j*_(*t*)], the shared environment, **Z**_*j*_(*t*), and the subpopulation parameters Θ_*N,j*_ = [***θ***_1,*j*_, ***θ***_2,*j*_, …, ***θ***_*N,j*_].

Similarly, the observation process (Equation (2)) becomes,

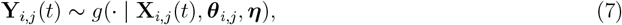

where **Y**_*i,j*_(*t*) is the fluorescence of the *i*th cell in the *j*th replicate. The snapshot data for the *j*th replicate at time *t > t*_0_ is *𝒴* _*j*_(*t*) = [**Y**_1,*j*_(*t*), **Y**_2,*j*_(*t*), …, **Y**_*N,j*_(*t*)]. We assume that the groups of *M* replicates are observed at the same *n* observations times *t*_1_ *< t*_2_ *<* … *< t*_*n*_, and denote *D*_*j*_ = [𝒴 _*j*_(*t*_1_), 𝒴_*j*_(*t*_2_), …, *𝒴*_*j*_ (*t*_*n*_)] for the series of snapshots nominally labelled as the *j*th replicate, as in reality there are *n × M* replicates as the process of taking the snapshot typically destroys the sample. This leads to the complete flow cytometry dataset, including the set of snapshots in replicate subgroups, to be represented by 𝒟= [*D*_1_, *D*_2_, …, *D*_*M*_].

Here, we assume that ***θ***_1,*j*_, ***θ***_2,*j*_, …, ***θ***_*N,j*_ are i.i.d. from a parametric distribution *p*(***θ*** | ***ϕ***_*j*_) with subgroup hyper-parameter, ***ϕ***_*j*_, representing the distribution parameters for the *j*th replicate. Across all replicates, these hyper-parameters ***ϕ***_1_, ***ϕ***_2_, …, ***ϕ***_*M*_, are distributed (i.i.d.) according to another parametric distribution, *p*(***ϕ*** | ***ψ***) where ***ψ*** are the between group hyper-parameters (Figure 2).

#### 2.3.2 Computational inference

To quantify both inter-experimental heterogeneity, characterised by *p*(· | ***ψ***), and intra-experimental heterogeneity, characterised by *p*_1_(***θ*** | ***ϕ***_1_), *p*_2_(***θ*** | ***ϕ***_2_), …, *p*_*M*_ (***θ*** | ***ϕ***_*M*_)., we aim to infer the hyper-parameters, ***ψ, ϕ***_1_, ***ϕ***_2_, … ***ϕ***_*M*_. That is,

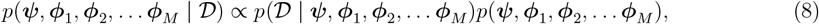

where the joint prior that enforces the hierarchical structure of the model is given by

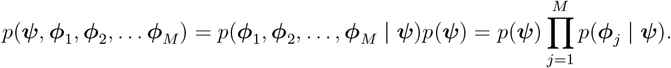

To perform parameter inference, we adopt an approximate Bayesian computation (ABC) approach [39–41]. That is, we approximate Equation (8) using the ABC posterior,

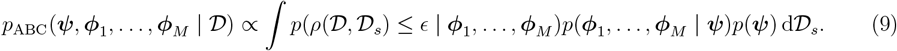

where 𝒟_*s*_ is simulated data conditional on the values of the specific parameters, ***ψ, ϕ***_1_, …, ***ϕ***_*M*_, and *ρ*(𝒟, *𝒟* ^*^) is a discrepancy metric with *ϵ >* 0 is a sufficiently small discrepancy threshold. Following Browning et al., [19] and Murphy et al., [32], we use a distribution matching discrepancy metric based on the Anderson-Darling distance. See the Appendix B.1 for details.

## 3 Results

To demonstrate the advantages of our approach, we perform simulations that represent *in silico* repeats of the experiments described in Section 2.1 based on published data [19, 25, 32]. The simulation setting enables experimental heterogeneity to be directly controlled and results to be compared to a known ground truth.

### 3.1 Nano-engineered particle-cell interaction experiments

Murphy et al., [32] consider an ODE model of particle-cell interactions (Section 2.2.2) with each cell, *i* = 1, 2, …, *N*, having its own particle association rate, *r*_*i*_,and carrying capacity, *K*_*i*_. In the pooled sample setting, they consider these parameters ***θ***_*i*_ = (*r*_*i*_, *K*_*i*_) to be log-Normally distributed,

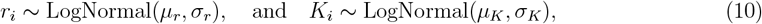

parameterised such that *µ*_*r*_ and *σ*_*r*_ are the mean and standard deviation of the association rate, and *µ*_*K*_ and *σ*_*K*_ are the mean and standard deviation of the carrying capacity. For inference, this leads to a vector of hyper-parameters of interest ***ϕ*** = (*µ*_*r*_, *σ*_*r*_, *µ*_*K*_, *σ*_*K*_).

#### 3.1.1 Synthetic data scenarios

Using the model in Section 2.2.2 and the parameter distributions (Equation (10)) for *r*_*i*_ and *K*_*i*_, we generate *in silico* data based on the experimental data collected by Faria et al., [25] as described in Section 2.1.1. We consider *M* = 3 experimental replicates each with *N* = 20,000 cells with simulated flow cytometry snapshots taken at *n* = 6 time points that are considered identically prepared technical replicates (*t*_1_ = 1 [hr], *t*_2_ = 2 [hr], *t*_3_ = 4 [hr], *t*_4_ = 8 [hr], *t*_5_ = 16 [hr], and *t*_6_ = 24 [hr]). Within this setting, we generate synthetic flow cytometry data for the following three scenarios:

- Scenario 1: All *M* experimental replicates are i.i.d. with hyper-parameters *µ*_*r*_ = 3.86125 *×* 10^−7^, *σ*_*r*_ = 4.56962 *×* 10^−7^, *µ*_*K*_ = 10, and *σ*_*K*_ = 2.
- Scenario 2: Cell parameters in experimental replicate 1 are generated with a larger mean association rate of *µ*_*r*_ = 9.0 *×* 10^−7^. Experimental replicates 2 and 3 are identical to Scenario 1.
- Scenario 3: Cell parameters in experimental replicate 1 are generated with a larger mean association rate *µ*_*r*_ = 9.0 *×* 10^−7^ and cell parameters in experimental replicate 2 are generated with a larger mean association rate of *µ*_*r*_ = 9.0 *×* 10^−7^ and a larger mean carrying capacity *µ*_*K*_ = 20. Experimental replicate 3 is identical to Scenario 1.

In all three scenarios we use, *c* = 1, *s* = 5.31 *×* 10^−5^, *v* = 1.01*×* 10^−6^, and *u*_0_ = 9.95 *×*10^7^. The specific values for these parameters are based on prior work of Murphy et al., [32].

We can consider Scenario 1 as the ideal experimental conditions for pooled sample analysis, whereas Scenarios 2 and 3 represent different ways the pooled assumptions may not hold. We not that the variation we have introduced to these parameters is fairly modest. Figure 3 provides an example of the synthetic flow cytometry data generated for Scenario 3. The larger mean association rate of replicate 1 (Figure 3(a)) leads to a subtle increase in the location of the peak fluorescence at early times, *t*≤ 4[hr], compared with replicate 3 (Figure 3(c), the basis of Scenario 1), though at later times, *t >* 8[hr], replicates 1 and 3 come into alignment. Replicate 2 generally is more diffuse than replicates 1 and 3 for all times, with the peak fluorescence approaching a higher steady state due to the higher mean carrying capacity (Figure 3(b)).

**Figure 3:**
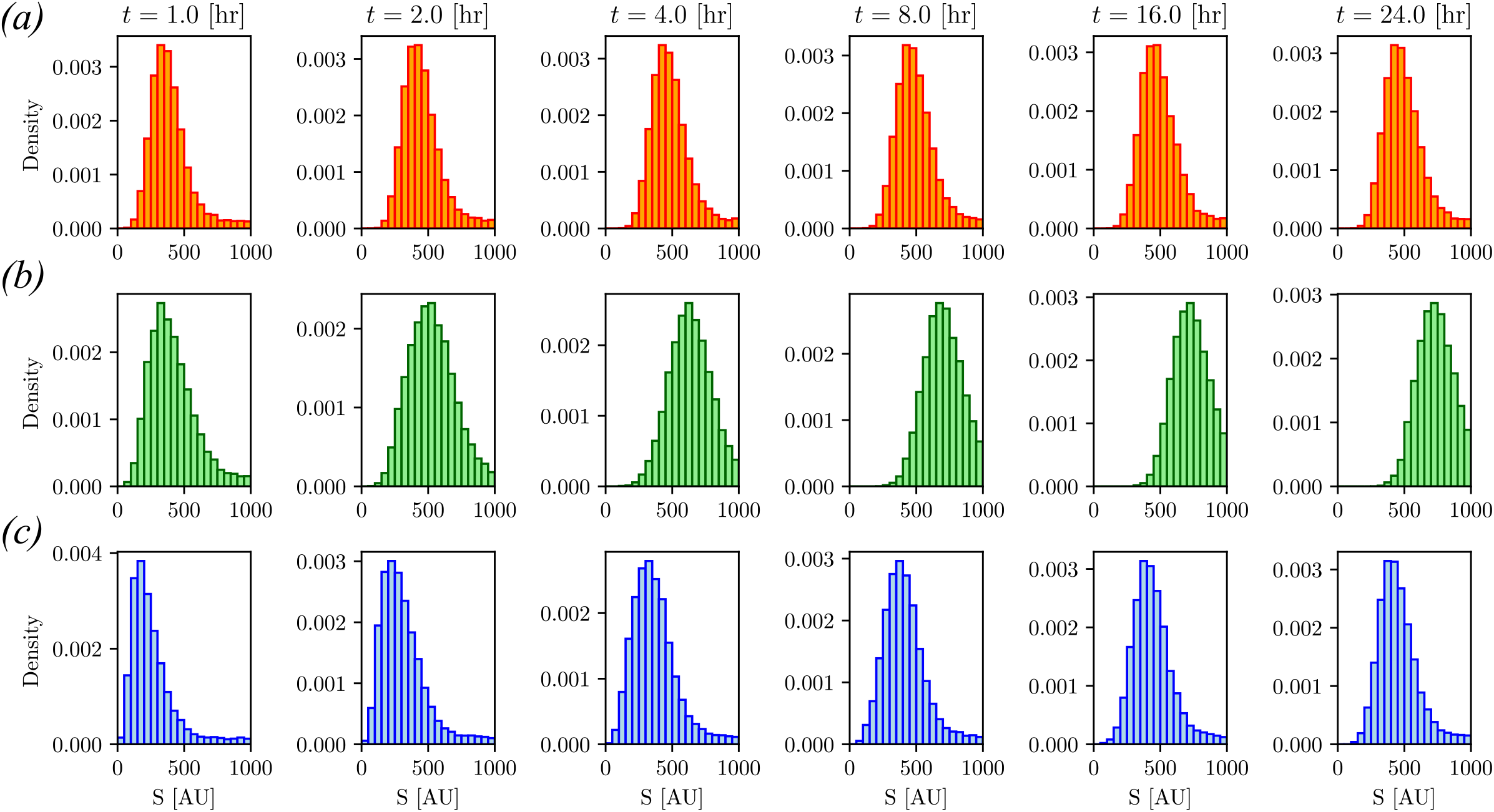
Synthetic flow cytometry data example simulating nano-particle cell interaction data using flow cytometry. Each histogram represents the simulated fluorescence *S* [AU] (that is Arbitrary Units) distribution for 20,000 cells interacting with nano-particles at one of six observation times (*t*_1_ = 1 [hr], *t*_2_ = 2 [hr], *t*_3_ = 4 [hr], *t*_4_ = 8 [hr], *t*_5_ = 16 [hr], and *t*_6_ = 24 [hr]). Replicate 1 (row (a)) is generated with a mean particle association rate of 9.0 *×* 10^−7^ and a mean carrying capacity of *µ*_*K*_ = 10. Replicate 2 (row (b)) is generated with a mean particle association rate of 3.86125 *×* 10^−7^ and a mean carrying capacity of *µ*_*K*_ = 20. Replicate 3 (row (c)) is generated with a mean particle association rate of 3.86125 *×* 10^−7^ and a mean carrying capacity of *µ*_*K*_ = 10. All replicates have the same standard deviation parameters for the association rate and carrying capacity of *σ*_*r*_ = 4.56962 *×* 10^−7^ and *σ*_*K*_ = 2, respectively.

#### 3.1.2 The effect of pooling on quantification of heterogeneity

Before presenting details of the pooled and hierarchical analysis in sections 3.1.3 and 3.1.4, we focus on estimated distributions of the individual cell parameters resulting from this analysis. In particular, we consider the heterogeneity in the particle association rate (Figure 4(a)) and carrying capacity (Figure 4(b)) that results from the analysis of synthetic data Scenario 3 (Section 3.1.1, Figure 3). This scenario captures a reasonable experimental setting for which the pooled i.i.d. assumptions are invalid due to experimental heterogeneity that is a confounding factor for the biological heterogeneity.

**Figure 4:**
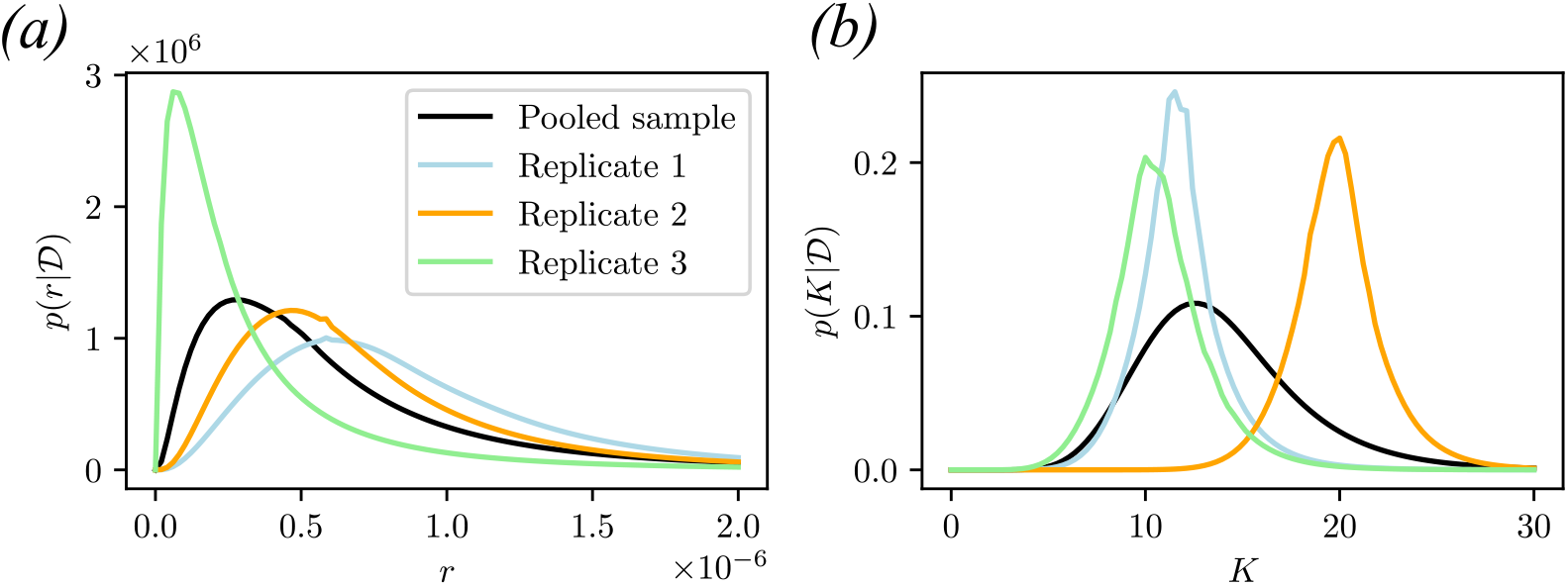
Heterogeneity in cell parameters resulting from pooled and hierarchical analysis of the particle-cell interaction model. Marginal distributions are shown for: (a) the particle association rate, *r*, and (b) the carrying capacity, *K*. Data Scenario 3 is used for this analysis (Section 3.1.1, Figure 3).

The pooled analysis adequately captures the variability in the cell parameter distributions with support that covers the bulk of each individual replicate distribution (Figure 4). However, the hierarchical analysis highlights that the actual biological heterogeneity is more complex with replicate 3 having a lower particle association rate (Figure 4(a)) and replicate 2 having larger carrying capacity (Figure 4(b)). These replicate specific distributions are not estimated independently, but rather, the hierarchical modelling structure allows for information sharing between replicates to provide quantification of experimental heterogeneity.

#### 3.1.3 Pooled sample analysis

We investigate the sensitivity of a pooled sample analysis to the i.i.d. assumptions. Following the pooled sample analysis protocol of Murphy et al., [32], we estimate the hyper-parameters, ***ϕ*** = (*µ*_*r*_, *σ*_*r*_, *µ*_*K*_, *σ*_*K*_), for each of the three synthetic data scenarios. These hyper-parameters characterise the heterogeneity (Equation (10)) in particle association rates and carrying capacities throughout the cell population, given the cell population dynamics under the model of Murphy et al., [32] (Section 2.2.2). The Bayesian analysis follows the approach described in Section 2.3, using independent uniform priors, *µ*_*r*_ ∼ 𝒰(0, 10^−6^), *µ*_*K*_ ∼ 𝒰(0, 50), *σ*_*r*_ ∼ 𝒰(0, 2 *×* 10^−6^), and *σ*_*K*_ ∼ 𝒰(0, 10). These priors are consistent with the analysis of Murphy et al., [32], and cover a range of physically viable values.

The results, as shown in Figure 5, indicate that the estimated hyper-parameters are sensitive to the synthetic data scenario. Firstly, the analysis of Scenario 1 results in posterior distributions that accurately recover the true hyper-parameters of *µ*_*r*_ = 3.86125 *×* 10^−7^ (Figure 5(a)), *µ*_*K*_ = 10 (Figure 5(b)), *σ*_*r*_ = 4.56962 *×* 10^−7^ (Figure 5(c)), and *σ*_*K*_ = 2 (Figure 5(d)). This is expected, since Scenario 1 represents our well-specified setting in which each replicate contains identical populations of cells. While this simulation validates the use of the pooled approach when when experimental replicates are genuinely the same [19, 32], the results for Scenarios 2 and 3 demonstrate substantial sensitivity of the posteriors given deviations in one of the replicates.

**Figure 5:**
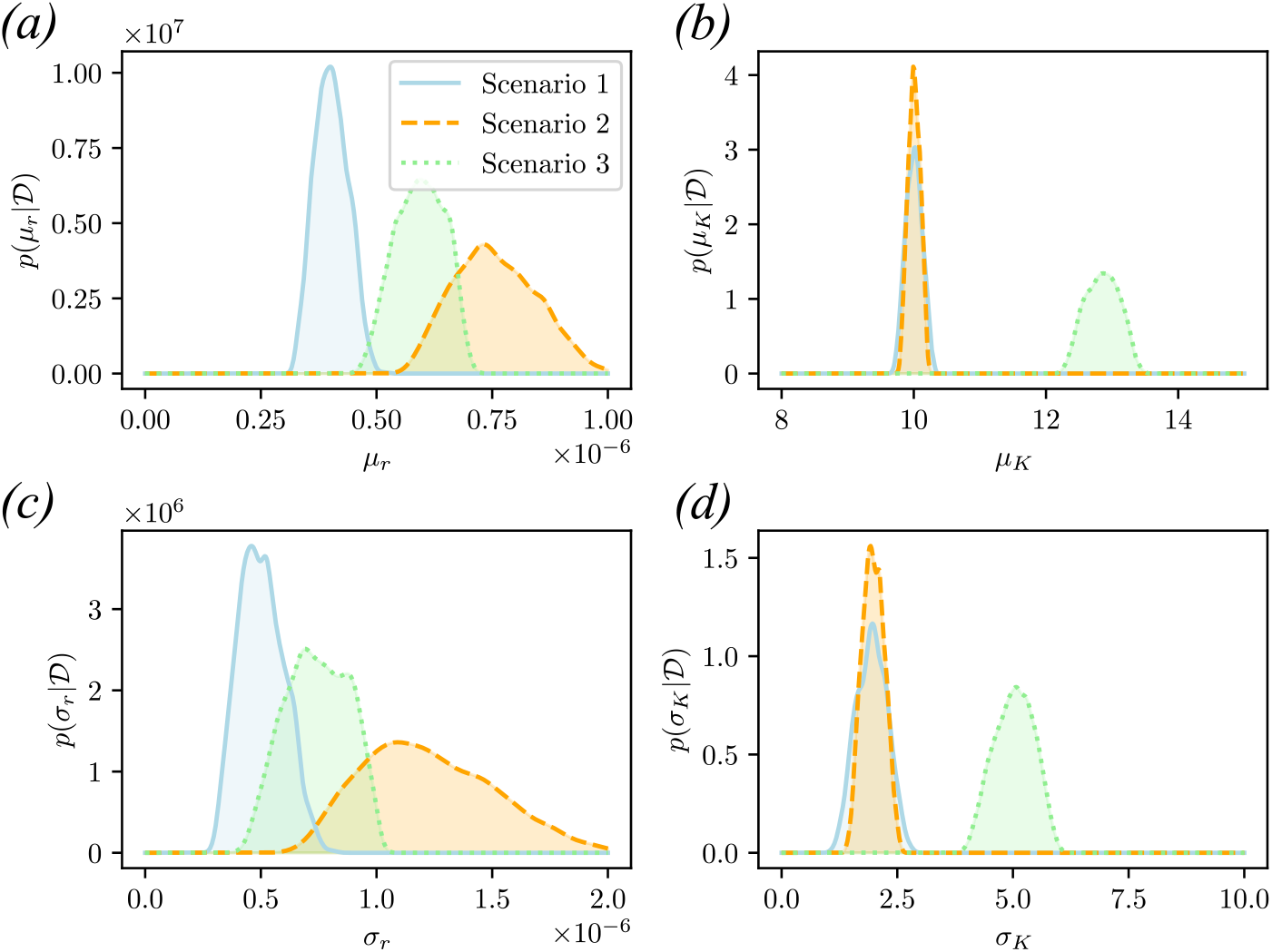
Pooled sample analysis of the nano-engineered particle-cell interaction experiment for the three synthetic data scenarios as described in Section 3.1.1. Marginal posterior distributions for the hyper-parameters: (a) mean association rate, *µ*_*r*_; (b) mean carrying capacity, *µ*_*K*_; (c) association rate standard deviation, *σ*_*r*_; and (d) carrying capacity standard deviation, *σ*_*K*_.

In Scenario 2, only one replicate was modified to have a mean association rate of *µ*_*r*_ = 9.0 *×* 10^−7^ resulting in potential outlier in one of the replicates (Figure 3(a)) compared with the unmodified replicates (Figure 3(c)). This outlier replicate strongly affects the estimates for both the pooled associate rate distributions with inflated means *µ*_*r*_ (Figures 5(a)) and standard deviations *σ*_*r*_ (Figures 5(c)). While the results do represent the best pooled sample inference given the outlier replicate, the results may not be meaningful for prediction in biological applications since the inferred heterogeneity does not represent any of the replicates accurately. The results are even more concerning for Scenario 3, where an addition replicate is modified to be an outlier association rate, *µ*_*r*_ = 9.0 *×* 10^−7^, and in carrying capacity *µ*_*K*_ = 20, since a shift in all four hyper-parameters occurs (Figure 5). This means that the heterogeneity estimates for the pooled population are not representative of any of the three replicate populations. These results motivate the adoption of our extended hierarchical model in experimental settings for which variation is expected between replicates.

#### 3.1.4 Hierarchical analysis

We now demonstrate the advantages of our hierarchical framework (Section 2.3) as a method for capturing heterogeneity in flow cytometry data when variation between replicates is present. We consider the same synthetic data scenarios (Section 3.1.1) and apply our hierarchical framework to the particle-cell interaction model (Section 2.2.2) of Murphy [32]. Specifically we consider the setting where the *i*th cell in the *j*th replicate population has its own associated rate, *r*_*i,j*_, and carrying capacity, *K*_*i,j*_. Following the pooled sample setting, we assume these parameters ***θ***_*i,j*_ = (*r*_*i,j*_, *K*_*i,j*_) are log-Normally distributed, however, we deviate from the pooled assumption by allowing each replicate population *j* = 1, 2, …, *M* to have its own mean association rate, *µ*_*r,j*_, and mean carrying capacity, *µ*_*K,j*_,

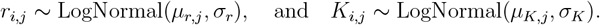

Note that the pooled sample model is recovered for the special case where *µ*_*r*,1_ = *µ*_*r*,2_ = … = *µ*_*r,M*_ and *µ*_*K*,1_ = *µ*_*K*,2_ = … = *µ*_*K,M*_. We further assign a Normal distribution to the replicate population means,

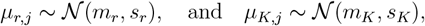

where *m*_*r*_ (resp. *m*_*K*_) and *s*_*r*_ (resp. *s*_*K*_) are the mean and standard deviation of the replicate population association rate (resp. carrying capacity) means. Thus we infer intra-experimental heterogeneity for replicate population *j* through inference of the hyper-parameters ***ϕ***_*j*_ = (*µ*_*r,j*_, *µ*_*K,j*_, *σ*_*r*_, *σ*_*K*_) and inter-experimental heterogeneity though the population level hyper-parameters, ***ψ*** = (*m*_*r*_, *m*_*K*_, *s*_*r*_, *s*_*K*_).

We perform Bayesian inference for the full joint posterior for the two levels of hyper-parameters (Equations (8) and (9)). For the population level parameters, we apply independent uniform priors for the means *m*_*r*_ ∼ 𝒰(0, 10^−6^), *m*_*K*_ ∼ 𝒰(0, 50), and weakly informative half Cauchy priors for the standard deviations *σ*_*r*_ ∼ Half-Cauchy(3 *×* 10^−7^) and *σ*_*k*_ ∼ Half-Cauchy(0.5). Note the shape parameters for the half Cauchy priors are determined based on the guidelines of Gelman et al., [42] with regard to selection of weakly informative priors for standard deviation terms in Bayesian hierarchical models with small numbers of groups.

The results, shown in Figure 6, highlight the value of our hierarchical approach. In Scenario 1 we accurately identify replicates are i.i.d. with *p*(*µ*_*r*,1_ | 𝒟) ≈ *p*(*µ*_*r*,2_ | 𝒟) ≈ *p*(*µ*_*r*,3_ | 𝒟) (Figure 6(a)–(c)), and *p*(*µ*_*K*,1_ | 𝒟) ≈ *p*(*µ*_*K*,2_ | 𝒟) ≈ *p*(*µ*_*K*,3_ | 𝒟) (Figure 6(e)–(g)). Furthermore the outlier replicates in Scenario 2 (replicate 1 with larger *µ*_*r*,1_; Figure 6(a)) and Scenario 3 (replicate 1 with larger *µ*_*r*,1_, and replicate 2 with larger *µ*_*K*,2_; Figure 6(a),(f)), are directly captured with only minor inflation in the variance of the posterior distributions for the remaining replicate populations (Figure 6(b),(c) for Scenario 2, and Figure 6(b),(c),(e),(g) for Scenario 3).

**Figure 6:**
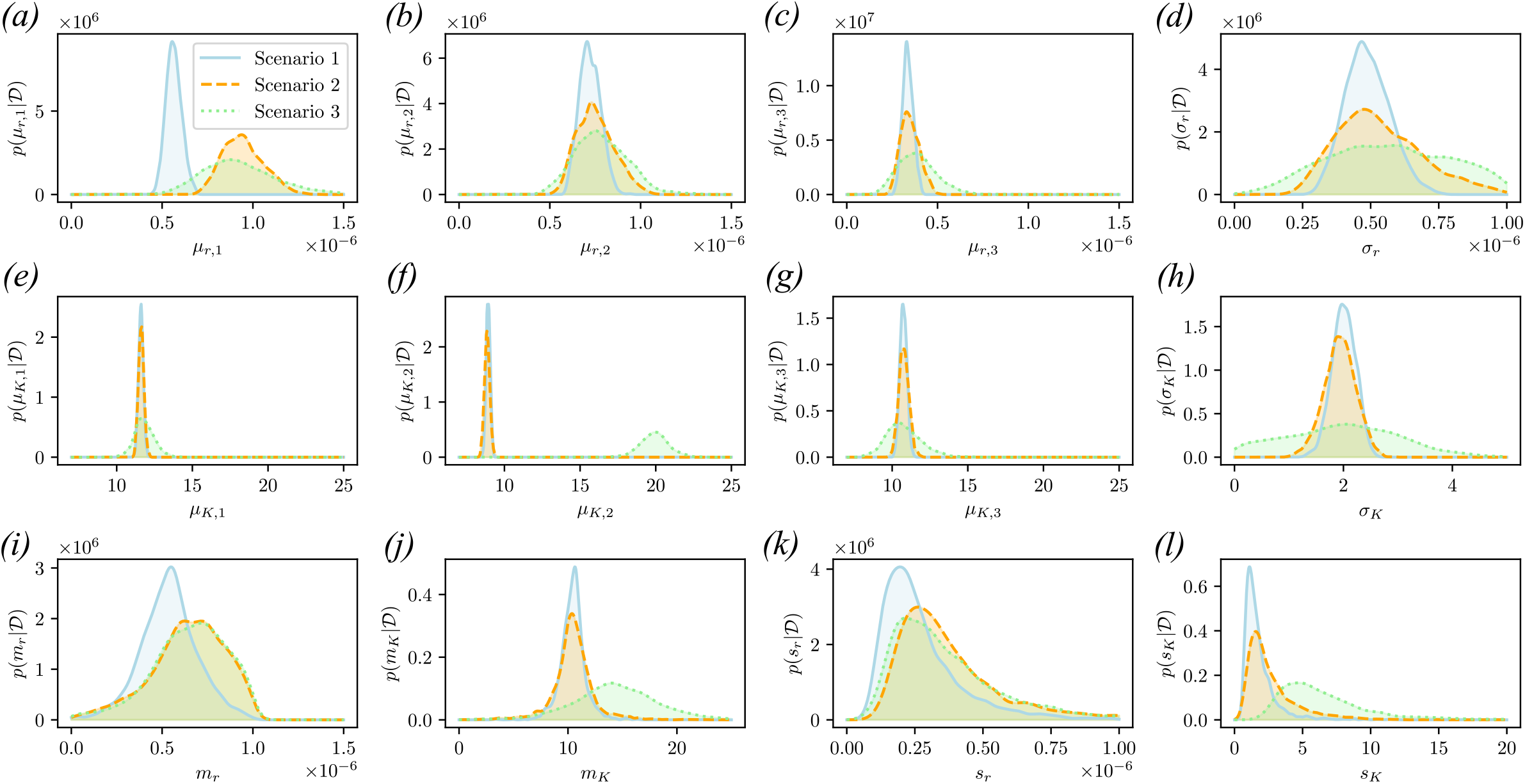
Hierarchical analysis of the nano-engineered particle-cell interaction experiment for the three synthetic data scenarios as described in Section 3.1.1. Marginal posterior distributions for the replicate hyper-parameters: (a)–(b) mean association rates, *µ*_*r,j*_ for replicates *j* = 1, 2, 3; (d) association rate standard deviation, *σ*_*r*_; (e)–(g) mean carrying capacity, *µ*_*K,j*_ for replicates *j* = 1, 2, 3; and (h) carrying capacity standard deviation, *σ*_*K*_. Marginal posterior distributions for the population level hyper-parameters: (i) mean of association rate means, *m*_*r*_; (j) mean of carrying capacity means, *m*_*K*_; (k) standard deviation of association rate means, *s*_*r*_; and (l) standard deviation of carrying capacity means, *s*_*K*_.

In addition to reliable characterisation of heterogeneity for each replicate population, we also obtain estimates of the inter-experimental heterogeneity that are insensitive to variation in a single replicate. This is evidenced by substantial overlap in the posterior probability densities around the true population level means (Figure 6(i),(j)) and standard deviations (Figure 6(k),(l)). In almost all cases, the true population level hyper-parameter is within the bulk density region of the posterior distribution. The exception, is the population standard deviation parameter for the carrying capacity mean *s*_*K*_ in Scenario 3 (Figure 6(l)), however, the deviation in the posterior distributions for *s*_*K*_ across scenarios is substantially less than that of the equivalent pooled sample analysis (Figure 5(d)). This robustness in population level inferences arises from the between-replicate information sharing inherent to the hierarchical model.

### 3.2 Dual-labelled probe internalisation experiments

To demonstrate the generality of or framework, we perform a similar simulation study based on the study of heterogeneity in cell internalisation processes by Browning et al., [19]. The internalisation process of a cell is modelled according an ODE system (Section 2.2.3) with each cell having its own internalisation rate, *λ*_*i*_, recycling rate, *β*_*i*_, and initial receptor density *R*_*i*_. The heterogeneity in these parameters is modelled, in the pooled sample setting, according to

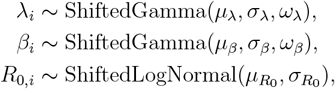

where *µ*_*λ*_, *σ*_*λ*_, and *ω*_*λ*_ (resp. *µ*_*β*_, *σ*_*β*_, and *ω*_*β*_) are the mean, standard deviation, and skewness of the internalisation rate (resp. recycling rate). Here, 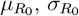 are the standard log-Normal parameters and the distribution of *R*_0,*i*_ is shifted such that 𝔼[*R*_0,*i*_] = 1 (See Browning et al., [19] for further details). For inference, these distribution hyper-parameters are of interest along with the disassociation rate *p*, that is, 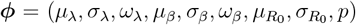. As per Section 2.2.3 (Equation (5)), parameters for to the observation process, ***η*** = (*α*_1_, *α*_2_, *σ*_1_, *σ*_2_), are also inferred, however, we do not focus on these here.

#### 3.2.1 Synthetic data scenarios

For the internalisation model (Section 2.2.3) we generate synthetic data inspired by the data collected by Browning et al., [19] as described in Section 2.1.2. Here, we have *M* = 3 replicates, each consisting of quenched/unquenched sample pairs of *N* = 1, 000 cells each. Simulated flow cytometry snapshots are taken at *n* = 7 observation times, *t*_1_ = 5 [min], *t*_2_ = 10 [min], *t*_3_ = 20 [min], *t*_4_ = 30 [min], *t*_5_ = 40 [min], *t*_6_ = 120 [min], and *t*_7_ = 180 [min]. As before (Section 3.1.1), we treat the different snapshots as being generated by identically prepared technical replicates, and thus having the same intra-experimental heterogeneity. We consider two scenarios based:

- Scenario 1: All *M* replicates are i.i.d., with hyper-parameters *µ*_*λ*_ = 0.08, *σ*_*λ*_ = 0.01, *ω*_*λ*_ = 0.9, *µ*_*β*_ = 0.05, *σ*_*β*_ = 0.005, *ω*_*β*_ = −1.0 *µ*_*R*_ = 0.05, and *σ*_*R*_ = 0.01.
- Scenario 2: Cell parameters in replicate 1 are generated with a smaller mean internalisation rate *µ*_*λ*_ = 0.02. Replicates 2 and 3 are identical to Scenario 1.

In both scenarios, *p* = 0.075, *α*_1_ = 8500, *α*_2_ = 38, *σ*_1_ = 5, and *σ*_2_ = 2. Parameter values are obtained based on the analysis of Browning et al., [19]. Figure 7 provides an example of the synthetic flow cytometry data generated for Scenario 2.

**Figure 7:**
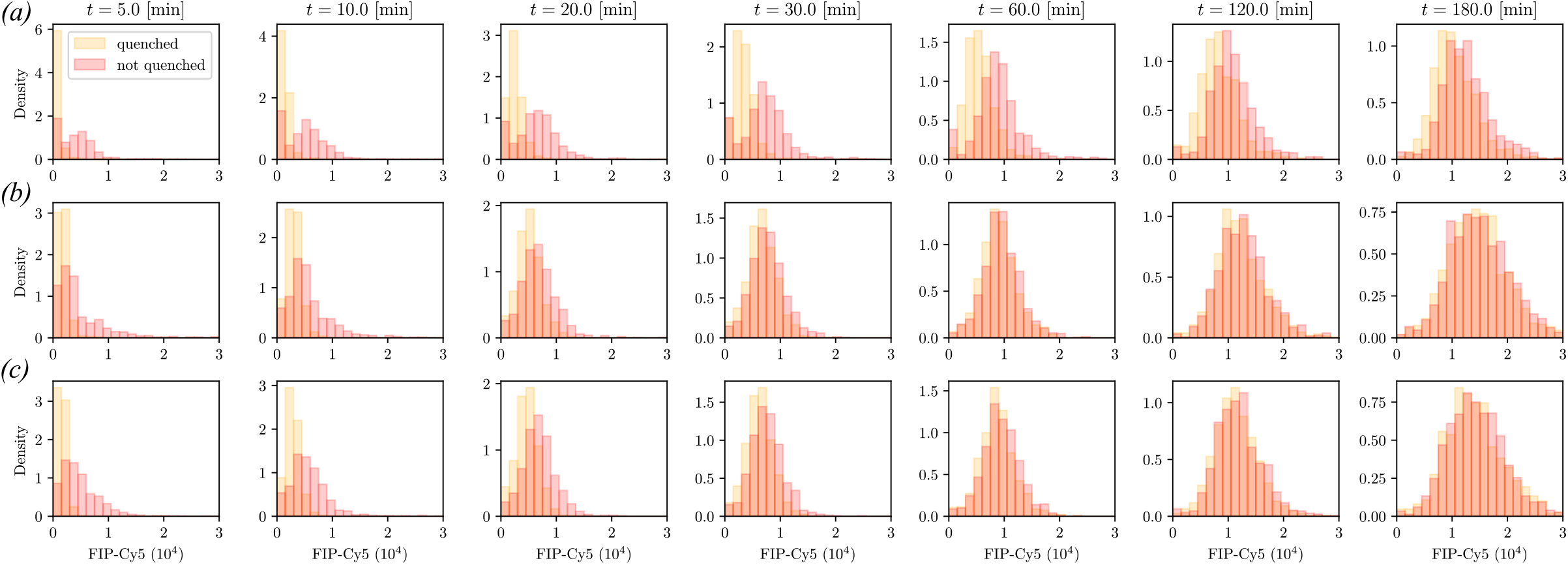
Synthetic flow cytometry data example simulating the dual-labelled probe internalisation experiments. Each histogram represent the simulated fluorescence distribution of the quenchable probe (FIP-Cy5) for 1, 000 cells in the quenched sample (orange) and 1, 000 cells in the unquenched sample (red) at one of seven observation times (*t*_1_ = 5 [min], *t*_2_ = 10 [min], *t*_3_ = 20 [min], *t*_4_ = 30 [min], *t*_5_ = 60 [min], *t*_6_ = 120 [min], and *t*_7_ = 180 [min]). Replicate 1 (row (a)) is generated with a mean internalisation rate of *µ*_*λ*_ = 0.02, a mean recycling rate of *µ*_*β*_ = 0.05 and a log-normal mean initial receptor concentration of *µ*_*R*_ = 0.05. Replicates 2 and 3 (row (b)-(c)) are generated with a mean internalisation rate of *µ*_*λ*_ = 0.08, a mean recycling rate of *µ*_*β*_ = 0.05 and a log-normal mean initial receptor concentration of *µ*_*R*_ = 0.05. All replicates have the same standard deviation parameters, *σ*_*λ*_ = 0.01, *σ*_*β*_ = 0.01, and *σ*_*R*_ = 0.01.

As with the nano-particle cell interaction example (Section 3.1.1), Scenario 1 represents the setting in which the pooled analysis is expected to be accurate, whereas Scenario 2 includes an outlier replicate (Figure 7(a)) with a smaller internalisation rate. This results in the quenched and unquenched samples being distinguishable throughout all snapshots. By comparison, replicates 2 and 3 lead to almost identical fluorescence distributions for quenched and unquenched at *t* = 30 [min].

#### 3.2.2 The effect of pooling on quantification of heterogeneity

In the pooled analysis, we note that the estimated variance of the internalisation rate is inflated to account for the variation across experimental replicates (Figure 8(a)). While the hierarchical approach provides more details with replicate specific parameter distributions, the inflated variance in the pooled approach seems an appropriate global approximation. However, the pooled analysis also inflates the variance of the recycling rate (Figure 8(b)) for which there is no substantial experimental heterogeneity. Conversely the pooled and hierarchical results for the initial receptor density are practically identical (Figure 8(c)). A reasonable explanation for the incorrect inflation of the *β* distribution is the relationship between *λ* and *β* in the model Equation (4) as receptor recycling is effectively a reversal of internalisation.

**Figure 8:**
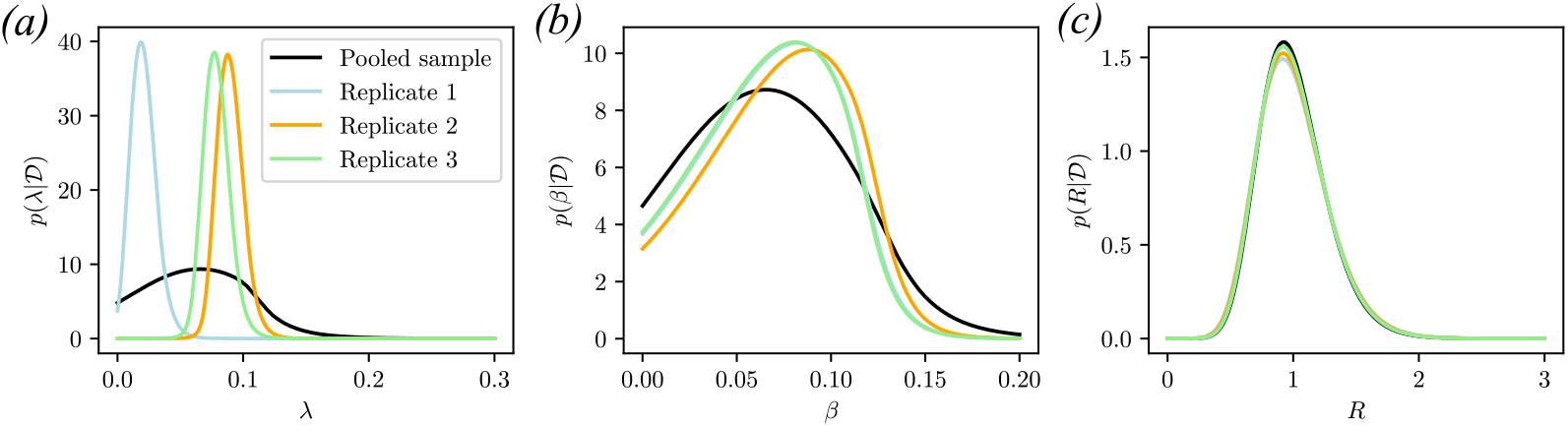
Heterogeneity in cell parameters resulting from pooled and hierarchical analysis of the internalisation model. Marginal distributions are shown for: (a) the internalisation rate, *λ*, (b) the recycling rate, *β*, and (c) the initial receptor density, *R*. Data Scenario 2 is used (Section 3.2.1, Figure 7) which exhibits variation across replicates in the mean internalisation rate.

#### 3.2.3 Pooled sample analysis

Here we investigate how deviations in the i.i.d. assumption affect quantification of biological heterogeneity, and the impact this has on estimates of global parameters, such as the disassociation probability *p*. For the pooled sample analysis we follow Browning et al., [19] to estimate the hyper-parameters, 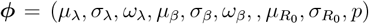 that relate to heterogeneity in internalisation rates, recycling rates, and the receptor concentrations. In addition, the purely chemical disassociation probability constant *p* is also.

The experimental heterogeneity incorporated into data Scenario 2 results in substantial estimated bias and variance inflation for both the mean (Figure 9(a)) and standard deviation (Figure 9(b)) parameters for the internalisation rate. Just as with the particle-cell interaction model (Section 3.1), the pooled sample analysis results are extremely sensitive to experimental heterogeneity. In addition, inferences of parameters with no biological or experimental heterogeneity, such as the disassociation rate *p* (Figure 9(c)), are impacted by experimental heterogeneity in the internalisation rate. Other parameter inferences are also inaccurate for Scenario 2, however, we focus here on the internalisation rate and the disassociation rate.

**Figure 9:**
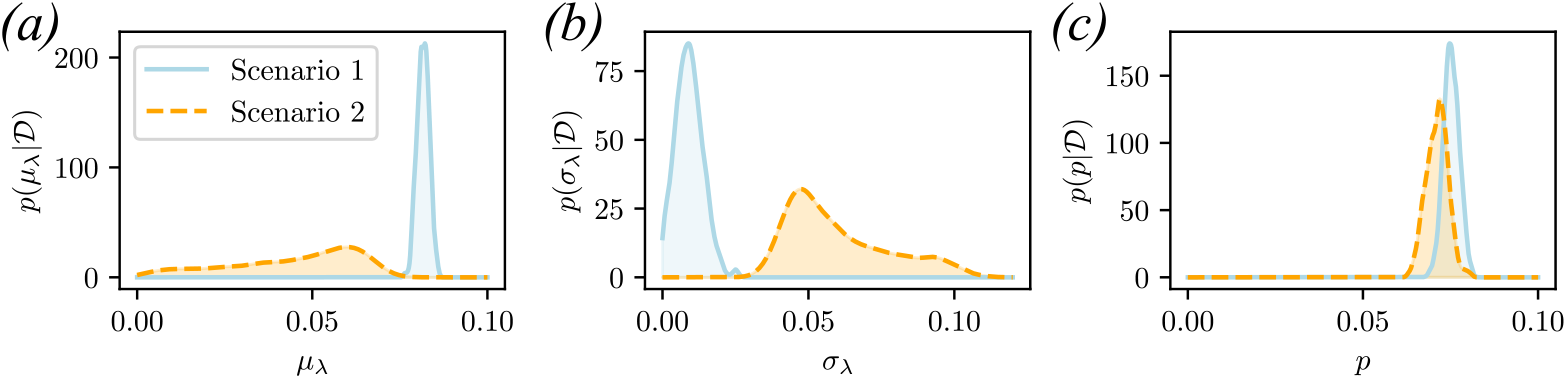
Pooled sample analysis of the dual-label probe internalisation experiment for the two synthetic data scenarios as described in Section 3.2.1. Marginal posterior distributions for the hyper-parameters: (a) mean internalisation rate, *µ*_*λ*_; (b) internalisation rate standard deviation, *σ*_*λ*_; and (d) the disassociation rate, *p*.

#### 3.2.4 Hierarchical analysis

Expanding the analysis of Browning et al., [19] according to our framework (Section 2.3) leads to the *i*th cell in the *j*th replicate population having its own internalisation rate, *λ*_*i,j*_, recycling rate, *β*_*i,j*_, and initial receptor density, *R*_*i,j*_. Further we allow the each of the replicate populations to have its own mean internalisation rate, *µ*_*λ,j*_, mean recycling rate, *µ*_*β,j*_, and mean initial receptor density, *µ*_*R,j*_,

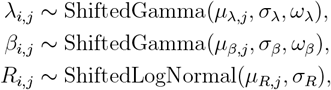

and treat the replicate population means as normally distributed,

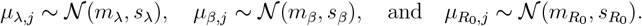

We observe improved results that accurately estimate the mean internalisation rate for each replicate population in both scenarios (Figure 10(a)–(c)) while quantifying the experimental heterogeneity across replicates (Figure 10(e)–(f)). Furthermore the disassociation rate is precisely the same in both scenarios ((Figure 10(g)). These results highlight that pooled analysis should only be used when there is high certainty in the i.i.d. assumption across replicates.

**Figure 10:**
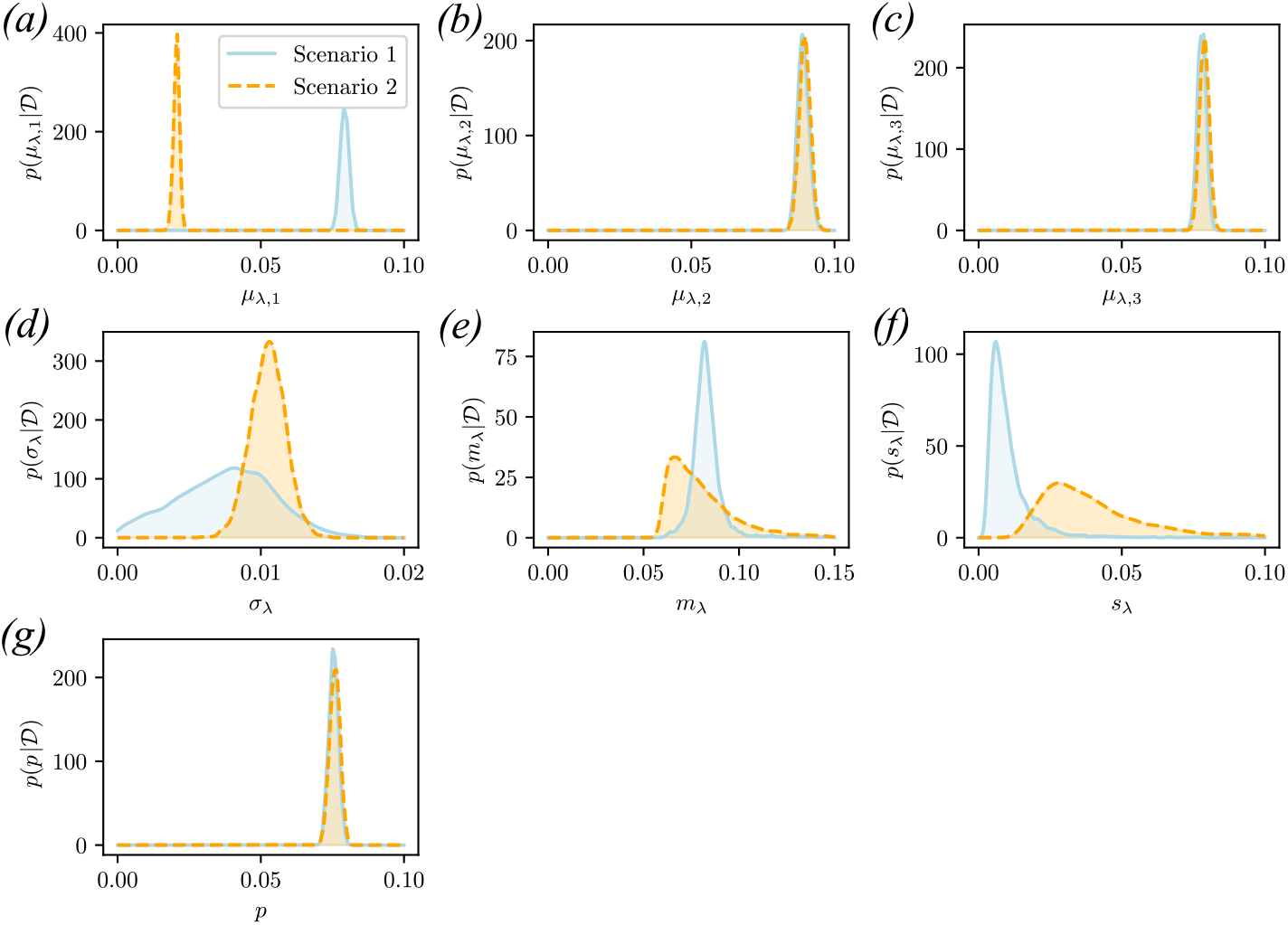
Hierarchical analysis of the dual-labelled probe internalisation experiment for the two synthetic data scenarios as described in Section 3.2.1. Marginal posterior distributions for the replicate hyper-parameters: (a)–(c) mean internalisation rates, *µ*_*λ,j*_ for replicates *j* = 1, 2, 3; (d) internalisation rate standard deviation, *σ*_*λ*_. Marginal posterior distributions for the population level hyper-parameters: (e) mean of internalisation rate means, *m*_*λ*_; (f) standard deviation of internalisation rate means, *s*_*λ*_; and (g) disassociation rate *p*.

## 4 Discussion

The reliable quantification of heterogeneity in cell populations is crucial to ensure accurate assessment and prediction of treatment efficacy [12–14], especially in the setting of targeted therapeutics [9–11]. Various approaches to modelling and analysis have been developed in the literature to quantify this heterogeneity using cell population snapshots obtained through flow cytometry. However, variability between experimental replicates is rarely considered. In this work, we present a Bayesian hierarchical framework to quantify biological and experimental heterogeneity simultaneously when using mechanistic models of cell dynamics. This is an important development, since previous approaches for quantification of heterogeneity using mechanistic models have only considered pooled samples and neglected experimental heterogeneity [19, 25, 26, 32].

We demonstrated our framework using a two-level hierarchical approach that modelled heterogeneity within experimental replicates and heterogeneity between each replicate. This could naturally be extended to include additional levels that might describe heterogeneity across different labs, instrument configurations, or cell lines. This would naturally come with additional computational challenges due to the larger dimensional parameter space, but such challenges are not insurmountable (Appendix A).

Here, we only consider inter-experimental heterogeneity in the parameter means across replicate populations. Experimental heterogeneity in the parameter variances are mathematically straightforward to introduce. However, some care should be taken to avoid non-identifiablity issues arising, especially for small numbers of replicates *M* [43]. Weakly informative priors, such as the Half Cauchy prior, are necessary for the variance hyper-parameter of the sub-population parameter means in the low replicate number setting *M* = 3 [42], and this would necessarily be more complex with experimental heterogeneity in variances [44]. As a result, we recommend that experimental heterogeneity in the replicate population parameter variances only be considered when the number of replicates, *M*, is large.

As our method was developed and presented as an extension to the pooled sample approaches [19, 32], we utilised similar modelling and analysis decisions. These could be adjusted as appropriate for other applications. For example, we assumed that heterogeneity is driven by variation between individual cells and that stochastic effects are negligible. In rare circumstances analytical solutions are available to account for non-negligible intrinsic noise [45], however, it is often necessary to use stochastic model, such as stochastic differential equations (SDEs), rather than deterministic ODEs [41, 46]. Since we rely on ABC for parameter inference, our framework is completely applicable to the SDE setting and other stochastic modelling approaches such as agent-based models.

The use of traditional simulation-based inference methods, such as ABC, could be considered a limitation due to the necessity of a tolerance threshold, *ϵ >* 0, that affects the accuracy of the posterior approximation [47–49]. In addition, targeting a very small *ϵ* is generally computationally burdensome and modern machine learning approaches are generally considered more efficient in this respect [49–52]. However, one desirable property of ABC, particularly in the context of biological modelling, is its robustness to model misspecification or model uncertainty [53–56]. While robust versions of modern machine learning methods continue to be developed [57–59], there is a serious lack of theoretical guarantees on how these methods behave in the misspecified setting. As a result, we still recommend ABC approaches when model misspecification is likely.

While our approach is developed with flow cytometry experiments in mind [19, 32], the method itself is general to any setting with population snapshot data taken from interacting individuals. For example, in the study of marine ecology, hierarchical Bayesian models are frequently used to describe coral reefs that vary across spatial regions [60, 61], however, mechanistic descriptions are typically only done on the level of individual reefs [62, 63]. Our approach would be well suited to account for spatial heterogeneity in these mechanistic models for more realistic large scale reef forecasts.

We have highlighted that caution must be taken when using pooled sample approaches to avoid potential bias or inaccuracies in quantification of biological heterogeneity. Our approach provides a means of reliable inference when experimental heterogeneity is present. Furthermore, we can quantify the heterogeneity across experiential replicates. This, in turn, could be used to identify potentially problematic replicates and improve experimental protocols. Due to these properties, our method has the potential to greatly enhance the biological insights that can be obtained through flow cytometry and advance developments in quantitative biology. Finally, our approach is generally applicable to other experimental techniques, such as microarrays that can result in drift and change between experimental replicates [64].

## Data availability

Data and Julia source code used in this study are publically available on Github.

## Authors’ contributions

DJW, APB, and MF conceived and designed the study. DJW, RJM, APB, STJ, and SAS provided analytical tools. SAS and DJW provided statistical tools. XZ, TPS, and DJW developed computational tools. XZ and DJW performed the research and drafted the article. All authors provided comments and approved the final version of the manuscript.

## Competing interests

The authors declare that we have no competing interests.

## Funding

This project was supported by the Australian Research Council (ARC). DJW is supported by an ARC Discovery Early Career Researcher Award (DE250100396). STJ and MF are supported by an ARC Discovery Project (DP230100380). SAS is supported by an ARC Discovery Project (DP260101134). DJW, APB, and STJ acknowledge support from the ARC Centre of Excellence for the Mathematical Analysis of Cellular Systems (MACSYS; CE230100001). APB thanks the Mathematical Institute, University of Oxford for a Hooke Research Fellowship. DJW acknowledges support from the Early Career Support Scheme and the Computational Bioimaging Group at Queensland University of Technology (QUT).

## Acknowledgements

The authors thank Adrianne Jenner and Adriana Zanca for helpful discussions. Computational resources where provided by the eResearch Office at QUT.

## A Extension to biological, technical, and experimental replicates

In the main manuscript, we consider a two layers of heterogeneity with details presented in Section 2.3. Here we will extend the approach to three levels heterogeneity that we will refer to as heterogeneity between *biological replicates, technical replicates*, and *experimental replicates*. Noting that terminology varies in the experimental biology literature, we will adopt the following terminology: each cell from a given cell line within a cell culture is a biological replicate; each cell culture population from a collection of identically prepared cell cultures is a technical replicate; and the reproduction of an entire experiment is an experimental replicate.

Suppose we adopt the random ODE approach from the main manuscript (Section 2.2.1) with *L* experiments each with *M* replicate cell populations of *N* cells. Let **X**_*i,j,k*_(*t*) and ***θ***_*i,j,k*_ be the state at time *t > t*_0_ and parameters of the *i*th cell (biological replicate) within the *j*th cell culture (technical replicate) from the *k*th experiment (experimental replicate). We have the system evolution and observation process as,

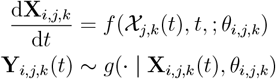

for *i* = 1, 2, …, *N, j* = 1, 2, …, *M*, and *k* = 1, 2, …, *L*. For ease of notation, we exclude the shared environment and observation process parameters described in the main manuscript. The state of the *j*th sub-population from the *k*th experiment is 𝒳_*j,k*_(*t*) = [**X**_1,*j,k*_(*t*), **X**_2,*j,k*_(*t*), …, **X**_*N,j,k*_(*t*)]. The snapshot data for the *j*th technical replicate from the *k*th experiment at time *t > t*_0_ is 𝒴_*j,k*_(*t*) = [**Y**_1,*j,k*_(*t*), **Y**_2,*j,k*_(*t*), … **Y**_*N,j,k*_(*t*)]. Then for *n* observation times, *t*_1_ *< t*_2_ *<* … *< t*_*n*_, the *j*th technical replicate from the *k* experiment is *D*_*j,k*_ = [𝒴_*j,k*_(*t*_1_), 𝒴_*j,k*_(*t*_2_), 𝒴_*j,k*_(*t*_*n*_)]. Then the *k*th experiment set of snapshots is 𝒟_*k*_ = [*D*_1,*k*_, *D*_2,*k*_, …, *D*_*M,k*_]. Finally the complete dataset, containing all biological, technical and experimental replicates is, 𝔇 = [𝒟_1_, 𝒟_2_, …, 𝒟_*L*_].

Biological heterogeneity is modelled by 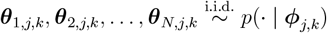 with each sub-population having its own population hyper-parameters ***ϕ***_*j,k*_. The heterogeneity in these hyper-parameters, that is technical heterogeneity, is modelled according to 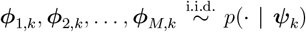 where ***ψ***_*k*_ is the experimental parameters for the *k*th experiment. Finally we have experimental heterogeneity given by 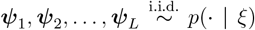 with *ξ* ∈ **Ξ** representing hyper-parameters charaterising the variation between experiments.

Just as is the main manuscript, estimating individual cell parameters ***θ***_*i,j,k*_ is considered meaningless, thus given data 𝔇 the aim is to infer the three levels of hyper parameters representing: experimental heterogeneity *ξ*; technical heterogeneity ***ψ***_*k*_ for *k* = 1, 2,, …, *L*; and biological heterogeneity, ***ϕ***_*j,k*_ for *j* = 1, 2, …, *M*, and *k* = 1, 2, …, *L*. This leads to the inference problem,

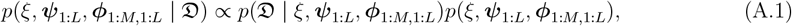

where ***ψ***_1:*L*_ = [***ψ***_1_, ***ψ***_2_, …, ***ψ***_*L*_], and ***ϕ***_1:*M*,1:*L*_ = [***ϕ***_1,1_, ***ϕ***_2,1_, …, ***ϕ***_*M*,1_, ***ϕ***_1,2_, ***ϕ***_2,2_, …, ***ϕ***_*M*,2_, …, ***ϕ***_*M,L*_]. The joint prior is,

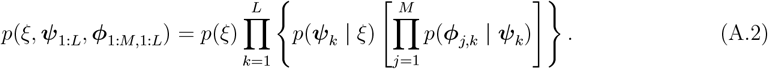

The likelihood is obtained through integrating out the individual cell parameters,

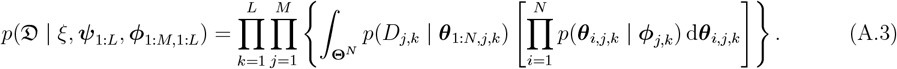

Standard Bayesian tools have be applied to sample from the posterior (Equation (A.1)) in the joint space **Φ**^*M*×*L*^ *×* **Ψ**^*L*^ *×* **Ξ**. However, the correlation structures could be quite complex and the likelihood evaluation (Equation (A.3)) could be challenging due to the *L × M* integrals over **Θ**^*N*^.

## B Computational inference

The two level hierarchical model is presented in the main manuscript (Section 2.3, Equation (8)) that captures heterogeneity both between replicates, characterised by *p*(· | ***ψ***), and within the cell subpopulation of each replicate, characterised by *p*_1_(· | ***ϕ***_1_), *p*_2_(· | ***ϕ***_2_), …, *p*_*M*_ (· | ***ϕ***_*M*_)., we aim to infer the hyper-parameters, ***ψ, ϕ***_1_, ***ϕ***_2_, … ***ϕ***_*M*_. That is,

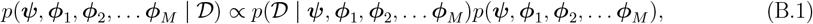

where the joint prior that enforces the hierarchical structure of the model is given by

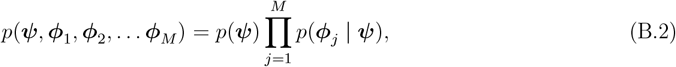

and the likelihood is

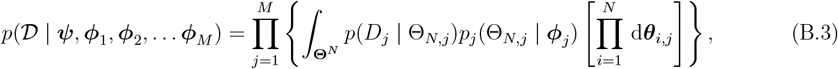

with 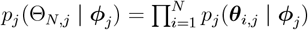.

In Equation (B.3) we integrate out the individual cell parameters (the ***θ***_*i,j*_’s) within the likelihood function. This leads to a complex likelihood evaluation while reducing the dimension of the inference problem substantially to the space of hyper-parameters **Φ**^*M*^ *×* **Ψ**. An alternative and equivalent formulation is to perform inference over the entire parameter space then perform the marginalisation over the full posterior, yielding

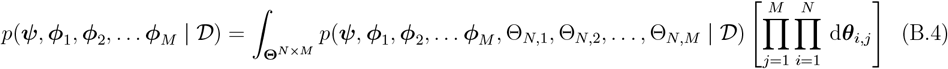

with

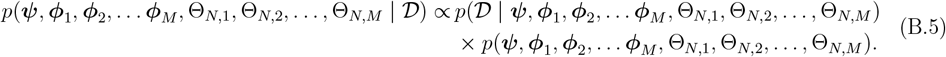

Here, we have the joint prior and likelihood,

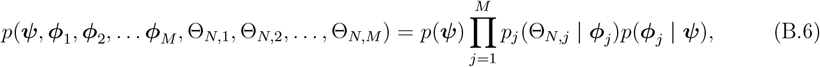

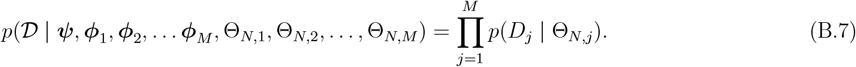

In this formulation, the likelihood function (Equation (B.7)) is relatively simple to evaluate (up to the complexity in obtaining a solution to the ODE system Equation (6)). However, it requires posterior sampling over the full space **Θ**^*N*×*M*^ *×* **Φ**^*M*^ *×* **Ψ**, which is challenging for standard Monte Carlo methods. To maintain the advantages of both formulations, that is, the lower dimensional inference problem from Equations (B.1)–(B.3) and the direct likelihood calculation from (B.4–(B.7)), we adopt an Approximate Bayesian computation (ABC) approach [39–41]. That is, we approximate Equation (B.1) using the ABC posterior,

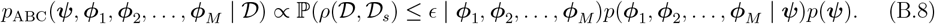

where 𝒟_*s*_ is simulated data and *ρ*(𝒟,𝒟_*s*_) is a distribution matching discrepancy metric based on the Anderson-Darling distance,

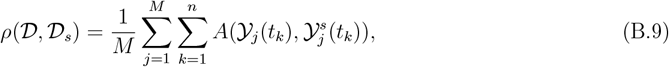

where 𝒴_*j*_(*t*_*k*_) = [*Y*_1,*j*_(*t*_*k*_), *Y*_2,*j*_(*t*_*k*_), …, *Y*_*N,j*_(*t*_*k*_)] is the real data snapshot for replicate *j* at time *t*_*k*_, and 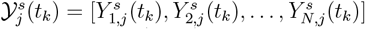 is the simulated data snapshot for replicate *j* at time *t*_*k*_. Here, the function *A*(𝒳, 𝒴) is the Anderson-Darling distance,

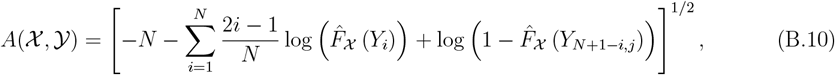

where 𝒳 = {*X*_1_, *X*_2_, …_*N*_ } and 𝒴 = {*Y*_1_, *Y*_2_, …, *Y*_*N*_ } are two sample sets and i s the empirical cumulative distribution for 𝒳, that is 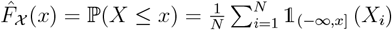.

We have that *p*_ABC_(*ϕ*_1_, *ϕ*_2_, …, *ϕ*_*M*_, *ψ* |𝒟) → *p*(*ψ, ϕ*_1_, *ϕ*_2_, … *ϕ*_*M*_ |𝒟) as *ϵ* → 0. The approach is intuitively encapsulated in the ABC rejection sampling scheme shown in algorithm B.1 for clarity, however, in practice it is not feasible to implement this method directly for small *ϵ* as required for accuracy. Instead we implement an adaptive ABC scheme based on sequential Monte Carlo methods of Drovandi and Pettitt [65] that refines *ϵ* through sequential importance resampling.

### Algorithm B.1

**ABC rejection sampling for hierarchical flow cytometry analysis.**

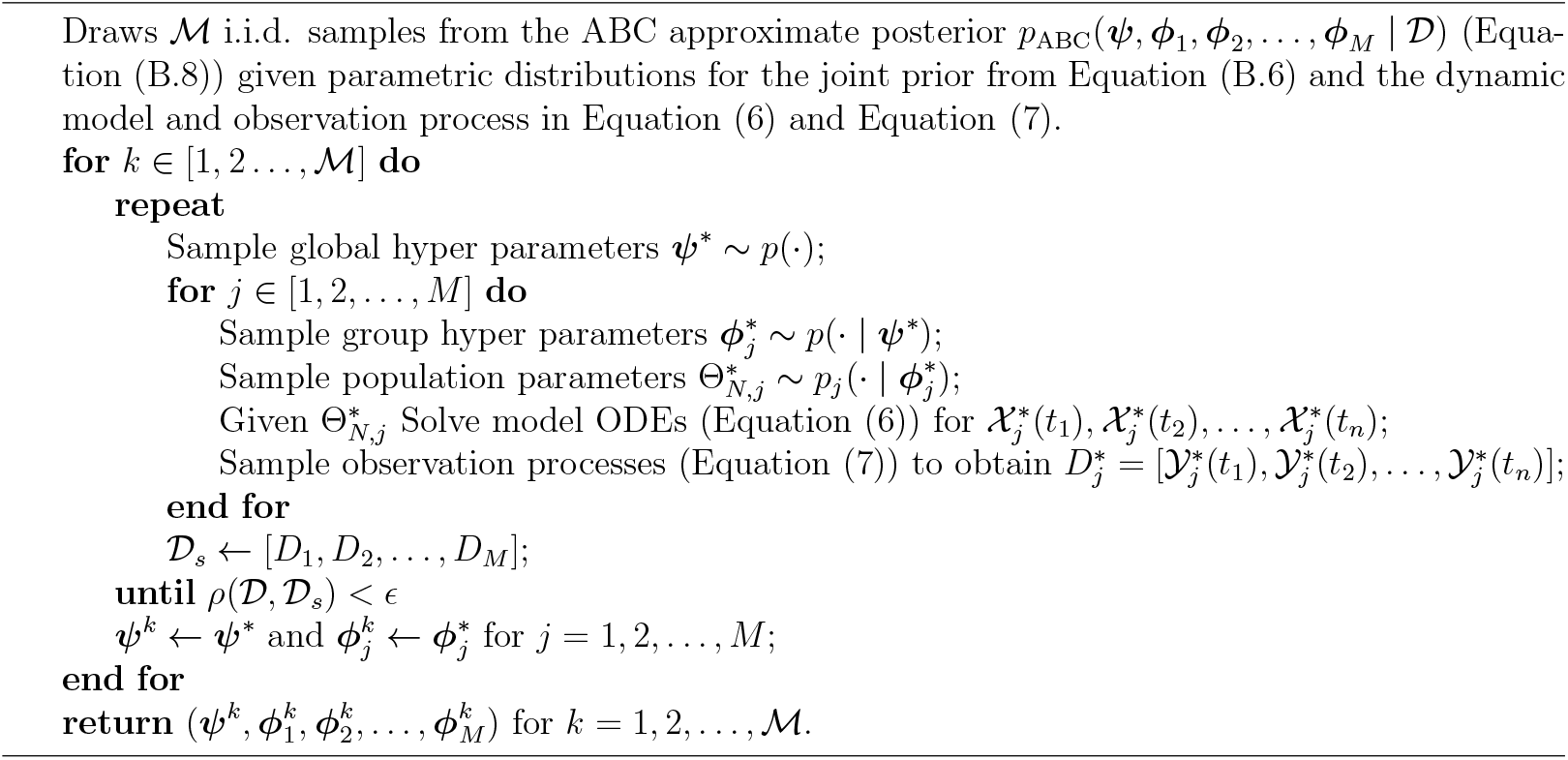

## Notes

### Competing Interest Statement

The authors have declared no competing interest.

